# Cancer risk and sexual conflict as constraints to body size evolution

**DOI:** 10.1101/2021.01.09.425980

**Authors:** E. Yagmur Erten, Hanna Kokko

## Abstract

Selection often favours large bodies, visible as Cope’s rule over macroevolutionary time — but size increases are not inevitable. One understudied cost of large bodies is the high number of cell divisions and the associated risk of oncogenic mutations. Our elasticity analysis shows that selection against a proportional increase in size becomes ever more intense with increasing body size if cancer is the sole selective agent. Thus cancer potentially halts body size increases even if no other constraint does so. We then provide multicellular realism with potentially sexually dimorphic body sizes and traits that control cell populations from zygote to maturity and beyond (ontogenetic management). This shows sexual conflict to extend to ontogeny; sexual dimorphism in mortalities and other life history measures may evolve even in the absence of any ecological causes underlying size- or sex-dependent mortality. Coadaptation of ontogenetic management and body size is required for substantial increases in size.

## Introduction

Body size varies greatly across the tree of life. Size is a key trait that correlates with numerous aspects of life history including longevity, age at maturity, and extrinsic mortality risk (Blueweiss et al., 1978; McCarthy et al., 2008; Healy et al., 2014). Selection can favour either small bodies (due to e.g. limited food availability, Blanckenhorn et al., 1995, or agility, Székely et al. 2004) or large ones (due to e.g. higher fecundity and mating success, Kingsolver and Pfennig, 2004, or resistance to starvation, Lindstedt and Boyce 1985). Examples for the latter are much more prevalent and the tendency of lineages to become larger is visible over macroevolutionary time (Stanley, 1973; Alroy, 1998; Heim et al., 2015) — a pattern so common that it has been given the name of a rule: Cope’s rule after Cope (1885) by Rensch (1948) (although its attribution to E. D. Cope’s work may be misplaced, see: Polly, 1998). Given that individual-level selection within a species can drive a macroevolutionary trend for lineages to increase in size (Kingsolver and Pfennig, 2004), why are not all organisms sized like whales and elephants — and why are these organisms themselves not even larger than they are? In reality, body size distribution is right-skewed, with few species occupying size categories close to the largest ones within a taxon (mammals: Gardezi and da Silva, 1999; Clauset and Erwin, 2008).

Resource limitation is a common constraint on life histories, and has clear potential to prevent open-ended selection for ever larger sizes (see Blanckenhorn, 2000, for a review of costs of a large size). In baleen whales, for instance, feeding efficiency increases with size, yet resource availability seems to ultimately prevent them from becoming even larger (Goldbogen et al., 2019). Taxon-specific reasons can also include physiological constraints associated with flight in birds (Tobalske, 2016) or oxygen transport in insects (Harrison et al., 2010). On a different level of selection, extinction risk can be elevated for large-bodied species that, all else being equal, form smaller populations (Damuth, 1981; Juanes, 1986; Clauset and Erwin, 2008); being large has been particularly detrimental to species once human impact started on this planet (Smith et al., 2018).

There is also an oncogenic and arguably underappreciated (but see: Galis and Metz, 2003; Kokko and Hochberg, 2015; Boddy et al., 2015; Sulak et al., 2016) constraint to becoming larger. Given that organisms vary much more in cell number than in cell size, achieving a larger size requires more cell divisions, assuming that the organisms in question all start from a single zygote. All else being equal, a higher number of total lifetime cell divisions increases the probability of an oncogenic mutation and translates into a potentially higher cancer risk (Peto, 1977). A potential counterargument to the importance of cancer as a constraint to body size evolution is Peto’s paradox, the observation that larger organisms do not appear to be more cancer prone than small ones in reality (Peto, 1977; Nunney, 1999). However, there is evidence that large-bodied lineages have had to evolve adaptations to reduce their cancer risk (Abegglen et al., 2015; Caulin et al., 2015; Sulak et al., 2016; Tollis et al., 2019; Martinez et al., 2020; Vazquez and Lynch, 2020). The problem may be solved in potentially lineage-specific ways, such as elephants refunctionalizing pre-existing pseudogenes (Vazquez et al., 2018). If this is the case, and if such solutions are not readily available at all times when selection would favour large size, it appears that the need to coevolve cancer defences with body size changes forms a potential constraint that can be overcome in some lineages but not in others. We are here interested in modelling this idea.

Based on previous models of multistage cancer process (e.g. Nunney, 1999; Calabrese and Shibata, 2010; Kokko and Hochberg, 2015; Webster, 2019), we first show how an approximation of finite lifespan allows to derive an analytical expression for selection to increase body size under cancer risk. The expression shows the generality of the finding that cancer risk selects against large bodies; specifically, should cancer be the sole selective agent, selection against a proportional increase in size becomes ever more intense with increasing body size. We thereafter move on to a simulation model, where we contrast cancer selecting for small bodies with reproductive success (potentially) selecting for large sizes. This model allows us to track body size evolution with and without simultaneous evolution of cancer defences, which we model in detail by tracking the health of the entire cell population for each individual. This model shows that large increases in size can only be realized if accompanied by the evolution of developmental strategies and cancer defences.

Our simulation model also allows us to incorporate an aspect of body size that is known to be of relevance in cancer biology (Fernandez and Morris, 2008; Boddy et al., 2015) as well as in life histories in general (Promislow, 1992; Tidière et al., 2015; Brooks and Garratt, 2017; Reedy et al., 2019): body size is very often a sexually dimorphic trait (Fairbairn et al., 2007). We take into account differences between the sexes by assuming that selection within a sex may take a different form in males than in females, and by allowing body size, cancer management strategies, or both to be sex-specific (or not). We show that sex-biased size expression allows sexes to diverge in their evolved body sizes and the consequent sexual size dimorphism results in sex differences in life histories and causes of mortality, with the larger sex suffering more from both developmental failures and cancer.

## Models

Cancer is typically modelled as a mutation accumulation process (Nunney, 1999; Calabrese and Shibata, 2010; Kokko and Hochberg, 2015), similarly to other multi-step failure models where organisms consist of ‘components’ that may fail over time (Gavrilov and Gavrilova, 2001; Laird and Sherratt, 2009; Webster, 2019). Our first model takes advantage of published results (as in: Nunney, 2018; Webster, 2019; Nunney, 2020), including a power function approximation for the risk of cancer (Webster, 2019). This approach enables us to derive the selection for body size increase assuming (1) cancer risk is body size dependent, (2) other aspects of life history (e.g. age at maturity, extrinsic mortality) do not scale with body size, (3) sexes are monomorphic (no sexual size dimorphism). We relax all these assumptions in the subsequent simulation model that allows us to include other sources of death, including extrinsic mortality, and to consider sex differences in body size under different reproductive scenarios.

### Analytical model

We define the probability that an individual has cancer by age *t* as:

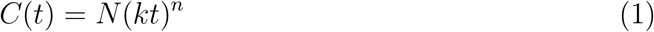

where *N* is the number of cell lineages under cancer risk, *k* is the rate of oncogenic mutations, *n* is the number of oncogenic steps (mutations within one cell lineage) until cancer occurs, and *t* is the age of the organism. This is a power law approximation (Webster, 2019) of a derivation that involves exponential decay functions (Nunney, 1999; Kokko and Hochberg, 2015); the approximation is very good until very old age, where the power law assumes all remaining individuals (which are few in number) to develop cancer while the exponential setting permits a very small proportion of individuals to escape the relevant oncogenic mutations for longer. We assume *N* to scale linearly with body size, hence we use it as a proxy for size and will hereafter refer to *N* simply as ‘(body) size’. The power law approximation in Eq.1 makes *C*(*t*) reach the value 1 at age *t*_max_ = *N^−^*^1*/n*^/*k*, which we take as the maximum age that can be reached. Irrespective of cancer, there is a constant death rate *μ* due to extrinsic causes (as in: Kokko and Hochberg, 2015; Boddy et al., 2015). We assume that the organism matures (instead of dying before reaching maturity) with probability *p*_mat_, and that deaths before maturity lead to zero fitness. Note that we keep our model parameters open in the sense of Cooper et al. (2018): we do not explicitly link high values of *μ*, or high age of maturity (defined below), to low *p*_mat_, even if (all else being equal) high extrinsic mortality operating for a long time would make maturation difficult; we allow fitness to be defined for any combination of these values and simply remark that real life may feature certain combinations more often than others.

After maturation, fitness (reproductive success) starts accruing at a constant rate until death, thus allowing us to calculate the lifetime fitness of an individual as:

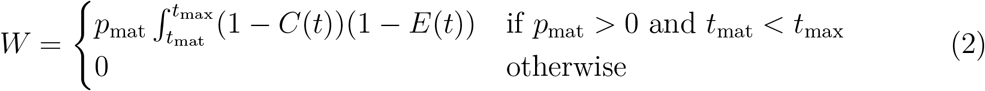

where *p*_mat_ is the probability of maturation, *t*_mat_ is the age at maturity, *E*(*t*) = 1 − *e^−μt^* captures the cumulative death probability from extrinsic mortality, and *μ* is the extrinsic mortality rate.

For analytical tractability, we here analyze the case where cancer is the sole death cause that can end an individual’s life (*μ* = 0), and refer the reader to the Supplementary Information for results regarding *μ* > 0. The first condition in Eq.2 permits positive fitness, and when it is satisfied, the expected fitness is:

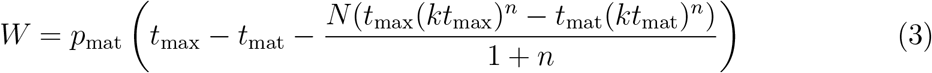

With *n* > 0 and *t*_max_ = *N^−^*^1*/n*^/*k*, the term *N* (*t*_max_(*kt*_max_)^*n*^ simplifies to *t*_max_, which decreases with body size *N*. Assuming neither maturation time nor probability scale with body size, we are ready derive our first result.

#### Cancer risk causes stronger selection against proportional body size change in larger organisms

Selection, *s*, is defined as the change in fitness *W* with body size *N*, but to compare selection across body sizes, it is important to note that a fixed absolute increase in body size represents a larger proportional body size change for a small than for a large organism. In other words, it is not meaningful to contrast the outcome for a 10 kg increase in weight across body sizes from a shrew to a whale. Therefore, to provide a meaningful comparison across several orders of magnitude of body sizes, we use elasticity (instead of a simple derivative 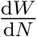) and define selection as 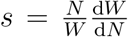. Elasticity calculates the proportional change in fitness when body size changes by a small proportion. Noting that *t*_max_ = *N^−^*^1*/n*^/*k* and assuming *n* > 0, *s* simplifies to:

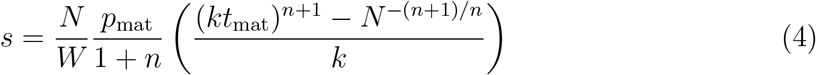

Eq.4 quantifies selection for a proportional change in body size, and if it is negative, selection favours smaller bodies. Since all parameters and *W* are positive, the sign of *s* (Eq.4) depends on the term (*kt*_mat_)^*n*+1^ − *N^−^*^(*n*+1)*/n*^. For the condition *t*_mat_ < *t*_max_ in Eq.2 to be satisfied, age at maturity needs to be smaller than maximum longevity, which implies *kt*_mat_ < *N^−^*^1*/n*^. For any *n* > 0, the term (*kt*_mat_)^*n*+1^ − *N^−^*^(*n*+1)*/n*^ will factorize to a positive term and the term (*kt*_mat_ − *N^−^*^1*/n*^). Since the latter term is negative, *s* will always fall below zero. In other words, selection against a proportional increase in body size occurs for all body sizes (an expected outcome since we assumed cancer risk to be the only life history feature in this model that responds to body size). Examples for three parameter combinations are shown in Fig.1).

**Figure 1:**
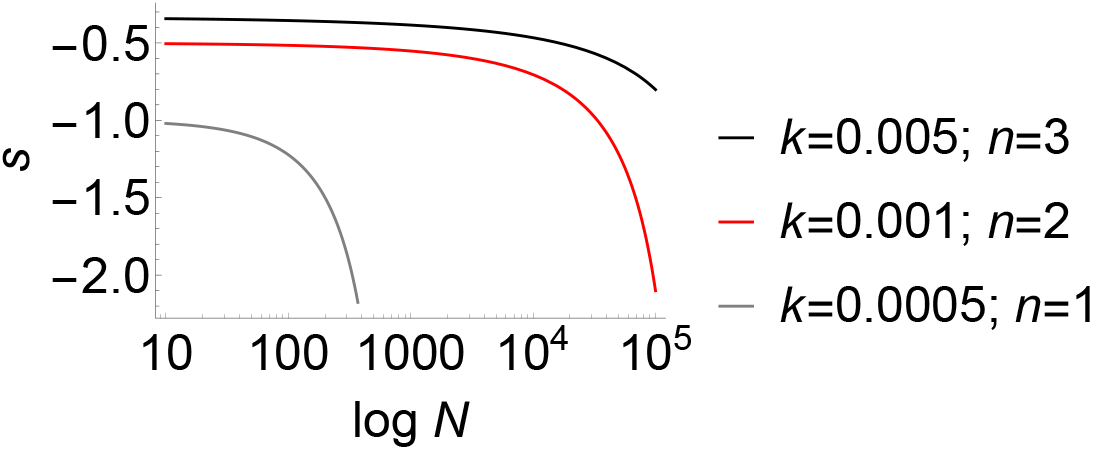
Selection *s*, defined as elasticity (proportional change in fitness with a proportional change in body size, *N*) according to Eq.4 plotted against log *N* and evaluated for *t*_mat_=2, *p*_mat_=1, and *k* and *n* as indicated in the figure legend. Fitness decreases with body size and the selection against a size increase is stronger in large organisms compared to the smaller ones.

Selection always acts against large body size, but to see how the strength of selection changes with body size, we can reorganize and simplify Eq.4 as:

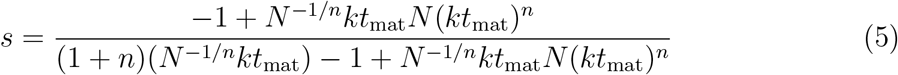

Multiplying both numerator and denominator with *N* ^1*/n*^ and assigning *A* = *kN* ^1*/n*^*t*_mat_, this can be written as:

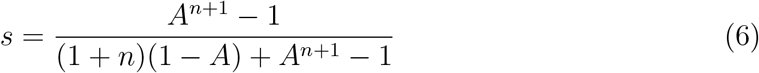

Since positive fitness is only possible if *t*_mat_ < *N^−^*^1*/n*^/*k*, we can deduce that *A* takes values between 0 and 1. Also, *A* increases with *N*, thus we can conclude that *s* not only always falls below zero, it also becomes more negative with increasing body size (Fig.1). In other words, there is a stronger selection against a proportional increase in size in large than in small organisms. Note that we derived this result with assuming that age at maturity or the probability of surviving to this age do not change with body size. If large mature size is difficult to reach, selection against large size would become stronger than indicated above. On the other hand, there are well documented benefits of being large (that do not relate to cancer) which are not included in this model; the Supplementary Information derives results based on the analytical model for such cases, and our simulation model (below) relaxes further assumptions.

### Simulation model

Our analysis thus far shows the potential of cancer risk to form a strong constraint that can prevent lineages from evolving large sizes, even in the absence of any other such constraint. Above, all actual selective benefits to be large were left outside the model (but see Supplementary Information). Accordingly, the model also does not comment on the potential sex-specific benefits of being larger than (same-sex) competitors and the consequent sex biases in outcomes such as cancer defences, body sizes, and realized incidence of cancer within an individual’s lifetime. In this section, we fill in these gaps by allowing (1) the relationship between size and fitness to differ between the sexes, and sex-biased expression of (2a) body size and (2b) cancer defences. The sex-dependence in assumptions can be switched on and off. For example, if (1) and (2a) are true but (2b) is not, females and males have to express the same cancer defences (ontogenetic management strategy, as defined below) due to a shared genetic architecture. This scenario allows us to investigate multiple potential outcomes, including but not limited to: cancer defences can become optimized for the larger sex, with the smaller sex being consequently ‘overprotected’ and dying rarely of cancer, or the defences can adapt to the smaller sex, with larger sex suffering from a high level of cancer incidence due to insufficient defences.

We also take advantage of our earlier study (Erten and Kokko, 2020) which produced candidate strategies for ontogenetic management, with clear potential to differ in how robustly they yield high fitness when expressed in larger (or smaller) bodies than before. Our earlier analysis had no sexes, and did not consider an actual evolutionary process towards a different body size (instead, each strategy was evolved under a fixed body size and thereafter tested in different-sized bodies, without considering whether selection can lead to the novel body size). In our current model, we take the strategies of Erten and Kokko (2020) (optimized for a fixed initial body size) as the ancestral population, and let evolution with sexual reproduction take its course in a population of *K* diploid individuals with even initial sex ratio. Each individual’s genotype consists of its ontogenetic management strategy Ω (modelled similarly to Erten and Kokko, 2020, see below), target mature tissue size *M*, and sex.

Each individual also has a phenotype comprising several traits. Some of the traits (sex, and the genetically determined ontogenetic management strategy) do not change as the individual ages, but may be sex-specific, as we permit sex-biased gene expression to a varying degree, depending on the scenario (see *Genotype and age-independent aspects of the phenotype*, below, for details). The ageing individual’s phenotype also has time-varying (age-dependent) components. Using the methods developed in Erten and Kokko (2020), we track the number and state of each individuals’ cells along three dimensions: how many cells are in each category, where categories are described by (i) number of divisions that have occurred since zygote state, (ii) level of differentiation reached, and (iii) level of damage (oncogenic steps completed; details in *Age-dependent aspects of the phenotype*, below).

#### Genotype and age-independent aspects of the phenotype

The ontogenetic management strategy, Ω, is defined phenotypically as in Erten and Kokko (2020), but has a more complex genetic basis than we assumed before. Briefly, it comprises seven traits involved in cellular differentiation and management and DNA damage response: probability of asymmetric cell divisions, *P* ; probability of differentiation in symmetric divisions, *Q*; maximum number of times a cell can divide, *H*; number of differentiation levels from a stem cell to a terminally-differentiated cell, *T* ; division propensity of a differentiation level compared to the previous one, *X*; DNA damage response threshold, *A*; and DNA damage response strength, *S*.

Each of the seven traits that comprise Ω and the additional trait of target body size *M* are based on three autosomal (diploid) unlinked loci. We indicate mid-parent values for each with subscripts f (for female-expressed), m (for male-expressed), and s (for shared, i.e. expressed in both sexes). We assume Mendelian inheritance and no parent-of-origin-specific expression. To permit sexually dimorphic trait expression, we introduce weight parameters *w_M_* and *w*_Ω_ (corresponding to variation in scenarios 2a and 2b, as listed above) that modulate the importance of the sex-specifically expressed loci versus the shared locus. For example, if *w*_Ω_ = 1, the ontogenetic strategy is constrained to be expressed identically in both sexes, and the phenotype Ω of either a female or a male (for each of the 7 traits) is the mean value of the two alleles at the shared locus Ω = Ω_s_. If *w*_Ω_ = 0, females and males express entirely different loci (while both sexes still inherit and carry all loci). In intermediate cases, e.g. *w*_Ω_ = 0.5, sexual dimorphism is permitted, but with genetic constraints making the phenotype of both sexes constrained to express values at the shared locus: Ω = 0.5Ω_s_ + 0.5Ω_f_ for females, and Ω = 0.5Ω_s_ + 0.5Ω_m_ for males.

#### Age-dependent aspects of phenotype

As described in Erten and Kokko (2020), individuals grow by dividing their cells according to their ontogenetic strategy Ω. Divisions alter the cell population of an individual in terms of how many cell divisions (since being a zygote) have occurred to produce a certain cell, how differentiated it is, and how much damage it has accumulated. We denote the number of cells that are terminally differentiated and can no longer divide as *N*_tissue_. At reaching its own target size *M*, such that *N*_tissue_ ≥ *M*, the individual is mature, and thereafter cell divisions only occur to replace cell loss such that tissue size is maintained at *M*.

Each time step gives individuals an opportunity to follow their ontogenetic strategy for growth (maximally one division per cell, should the strategy dictate divisions are needed, i.e. the organism is below its target size). Divisions can occur in cells that are not terminally-differentiated (‘stemlike’ cells) and move those cells towards a more frequently divided (according to *X*), more differentiated (according to *P* and *Q*), and — probabilistically — more damaged level, until terminal differentiation, as described in Erten and Kokko (2020). Both stemlike and terminally-differentiated cells can die due to baseline cellular mortality (at a rate *ν*), due to having reached their maximum number of divisions (the Hayflick limit, coded as the trait *H*), or as a DNA damage response (depending on *A* and *S*). At the end of each time step, the entire individual may die.

We track organismal deaths from five causes (details given in Erten and Kokko, 2020): 1) extrinsic mortality (with constant rate *μ* applied both to pre- and post-maturity), 2) depleting their stemlike cells (which we divide into pre-maturation and post-maturation), 3) losing all the cells before maturity, 4) cancer, and 5) falling below a threshold of tissue cells needed for somatic maintenance. Cause 1 occurs randomly (irrespective of body size or other traits), and is applied if a random number falls below *μ* at a given time step. Causes 2 to 5 are dependent on the health of the entire cell population of the individual: cause 2 happens if the organism does not have any stemlike cells left, cause 3 happens if cell death events remove all the cells of the organism (both stemlike and tissue) at a given time step, cause 4 occurs when at least one cell of the organism reaches to the highest level of oncogenic damage, and finally cause 5 implies insufficient somatic maintenance and occurs when the number of tissue cells falls below a threshold (set to 80% of the organism’s target mature size) after maturation.

#### Reproduction

We keep the population at a constant size, using *K* to denote the carrying capacity. If a time step involved one or more deaths of individuals, the formed vacancies will be filled by a new offspring (if there are mature individuals who can reproduce; otherwise the population is let temporarily fall below *K*). Among-female competition determines the choice of the mother who generates this offspring, likewise males compete among themselves for the status of the sire. Only individuals who have reached their mature size can participate in female-female or male-male competition.

An individual’s probability of winning the reproductive competition (being successful at placing a new offspring into the population; hereafter ‘winning’ for brevity) is proportional to its current competitive ability, which we define in two alternative ways. In the case of size-dependent selection, competitive ability is determined by tissue size *N*_tissue_, while in the case of somatic budget selection, competitive ability is *B*(*t*) at the time step *t* when competition occurs, defined as:

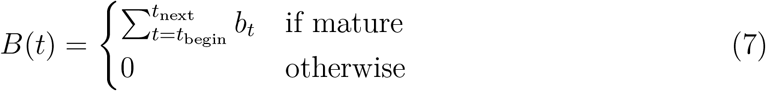

where *b_t_* is the ratio of tissue cells, *N*_tissue_, to the total body size (*N*_total_, which includes both the tissue cells and the dividable stemlike cells), such that *b_t_* = *N*_tissue_*/N*_total_ at time *t*. We assume that the budget *B*(*t*), simply referred to as *B* hereafter, starts to accumulate once the organism reaches its target mature size and the entire budget is spent when the organism reproduces. Therefore, *t*_begin_ refers to either maturation or the end of the previous reproduction, whereas *t*_next_ is the time step at which the organism is evaluated for reproduction.

All mature individuals can participate in reproductive competition whenever vacancies arise. Those who succeed in placing one or more offspring into the population lose their current budget as reproductive effort, and begin accumulating it again thereafter. Competitors who did not win keep their budget intact, i.e. we assume actual reproduction, rather than competing to do so, is costly.

For sex differences, we use four different assumption sets (scenarios) to model reproductive competition: success depends on 1) size *N*_tissue_ or 2) reproductive budget *B* in both males and females 3) males are picked for *N*_tissue_ and females for *B*, 4) males are picked for *B* and females for *N*_tissue_. In each case, the propensity of being chosen is proportional to the relevant competitive trait (either *N*_tissue_ or *B*, depending on scenario). Note that for size-dependent selection, we only count differentiated tissues to contribute to fitness, to prevent undifferentiated cell masses from having good reproductive success: a successful ontogenetic management strategy has to produce organisms with functional organs.

Inheritance of all traits is Mendelian, sex ratio at birth is 1:1, and individuals begin their lives with age set to 0. Each locus is inherited with a mutation in either the mother’s or the father’s gamete, such that in each generation one allele has a mutation while the other is inherited faithfully from the mother or father. Mutation size (*δ*) is normally distributed with *δ* ~ *N* (0, 0.5), and change allelic values as follows, exemplified for the maternally-inherited allele of the female-expressed target mature size *M*_f,mother_: the new allelic value is 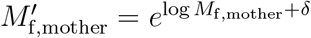 where *δ* ~ *N* (0, 0.5); similarly for all other cases (shared and male-expressed). Ω mutates like *M*, but for consistency with earlier work, we do not allow the novel allelic values to go outside of the range we explored in Erten and Kokko (2020). We optionally make the ontogenetic strategy a non-evolvable trait by disallowing mutations in it.

#### Simulation runs

We begin each simulation run with a population of *K* = 1000 individuals (500 males and 500 females) all sharing the same ontogenetic strategy Ω and target body size *M*. Since completely random Ω are unlikely to yield viable bodies, we choose four different ancestral populations from those that evolved under fixed body size in Erten and Kokko (2020). This choice allows us to bypass the initial search for viable ontogenies, but the outcomes reported in Erten and Kokko (2020), being independent of sex (and lacking the comparison between size-based and budget-based selection), are not necessarily final for evolution in a sex-specific setting that is our current focus of interest. We denote the initial ontogenetic strategies as Ω_*i*_, where *i* = 1, 2, 3, 4 indicates distinct strategies used in our simulations.

We introduce variation in the evolvable traits (target body size as default; also Ω depending on the scenario) by allowing each individual in the initial population to start with slightly different trait values, randomly shifted from the initial ones following the mutation process explained above (in the section *Reprodcution*), with *δ* ~ *N* (0, 0.05). When Ω is kept constant (i.e. non-evolvable), we introduce no variation in Ω and assume that the population is monomorphic with respect to Ω and initialise all individuals with previously-optimized ontogenetic management strategies.

A time step starts with cells of each individual dividing, differentiating and potentially dying (following their ontogenetic strategy, as described above). Entire individuals can then die, due to one of the causes listed in the section *Age-dependent aspects of phenotype*. Those who survived have their age increased by one time step (i.e. d*t*=1), and their current body size, and overall budget *B* updated. The removal of dead individuals creates vacancies that are then filled with the offspring of surviving mature individuals who compete for reproduction as described above. A time step ends with the addition of new offspring (if any) to the population, taking into account their mutations; offspring start with just 1 cell (the zygote).

We run the simulations for 10000 time steps (unless indicated otherwise); we use a fixed stopping time for computational reasons as by this time it proved clear whether scenarios lead to an asymptotic size quickly or permit more open-ended growth (note that our model is not merely individual-based but also tracks each individual’s cell population along three different axes of cell state — number of divisions completed, differentiation level, and oncogenic damage accumulated — making it extremely computationally intensive).

### Results of the sexual model

#### Body size increases under size selection

Overall, when reproductive competition was settled based on body size in both sexes, *N*_tissue_, size at maturity increased (Fig.2a-e). However, the magnitude of the increase was strongly dependent on whether ontogeny management (Ω) was allowed to coevolve with body size. If it was not, size increases remained modest (Fig.2a-b), while ontogenetic coevolution permitted sizes to increase over several orders of magnitude in a potentially open-ended fashion (Fig.2c-e). In other words, coevolution of ontogeny management was imperative for large increases in size.

**Figure 2:**
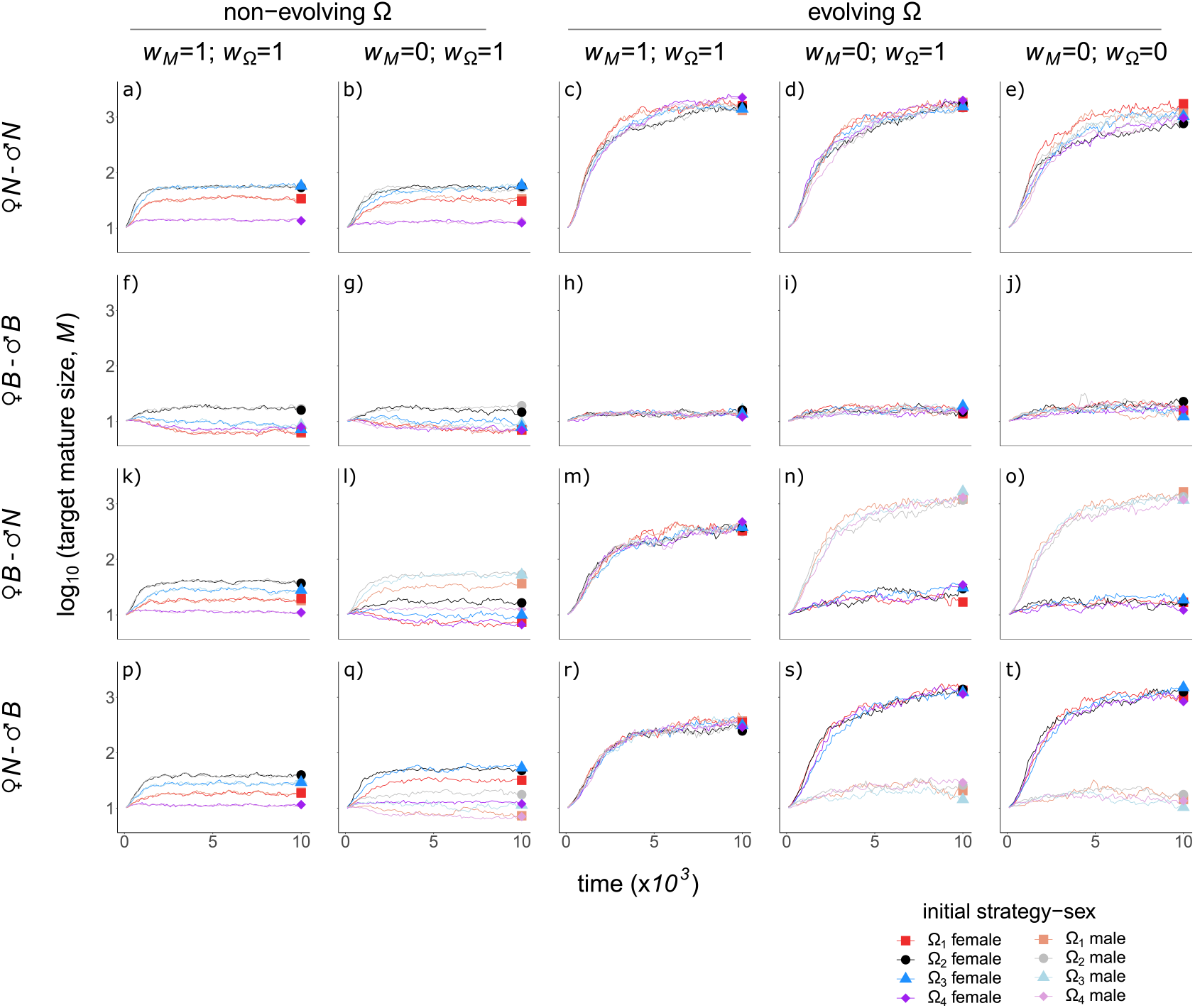
Body size evolution. Columns show whether ontogeny management is evolvable, sex-specificity of body size (*w_M_*) and ontogeny management (*w*_Ω_; equal to 1 when sexes express shared alleles and 0 if expression is sex-specific) as indicated. Rows depict reproductive scenarios, top to bottom: size selection in both sexes (*N*_tissue_ shortened as *N*, *N*-*N*), budget selection in both sexes (*B*-*B*), budget-selection in females and size-selection in males (*B*-*N*), size-selection in females and budget-selection in males (*N*-*B*). In each panel, x-axis: time steps in the simulation; y-axis: mean target tissue size in logarithmic scale (log_10_ *M*) for the mature individuals in the population. Colours and dots show the initial strategy and sex as indicated. When both sexes were size-selected, target mature size increased particularly when ontogeny management coevolved with body size, whereas budget selection resulted in small-bodied lineages. Sex differences in reproductive scenarios led to either size increases in both sexes (when *w_M_* =1) or sexual size dimorphism (when *w_M_* =0). All simulations were run for extrinsic mortality *μ*=0.01, cell turnover rate *ν*=0.0001, oncogenic mutation rate *c*=0.01, number of oncogenic steps *n*=3. See Erten and Kokko (2020) for details. Results shown for mature individuals in the population for intervals of 100 time steps.

#### Budget selection favours small body sizes

When optimal management of somatic budget (*B*) is used as a determinant of reproductive success for both sexes, body sizes remained small across all the scenarios (Fig.2f-j). This included cases (three out of four lineages) with body size shrinkage when body size was an evolvable trait but ontogeny management was not (Fig.2f-g). Our earlier results suggest that stem cell management is more challenging in large bodies (Erten and Kokko, 2020) and this, combined with a higher age at maturity in larger organisms (Fig.3b), can favour small-sized organisms under budget selection (similarly to Erten and Kokko, 2020, where fitness, *W* = *N*_tissue_*/N*_total_, was shown to decrease with target mature size). We did not, however, put a straightforward selection coefficient for smallness per se, and indeed, allowing coevolution with the ontogeny management strategy increased body sizes slightly in both sexes (Fig.2h-j).

**Figure 3:**
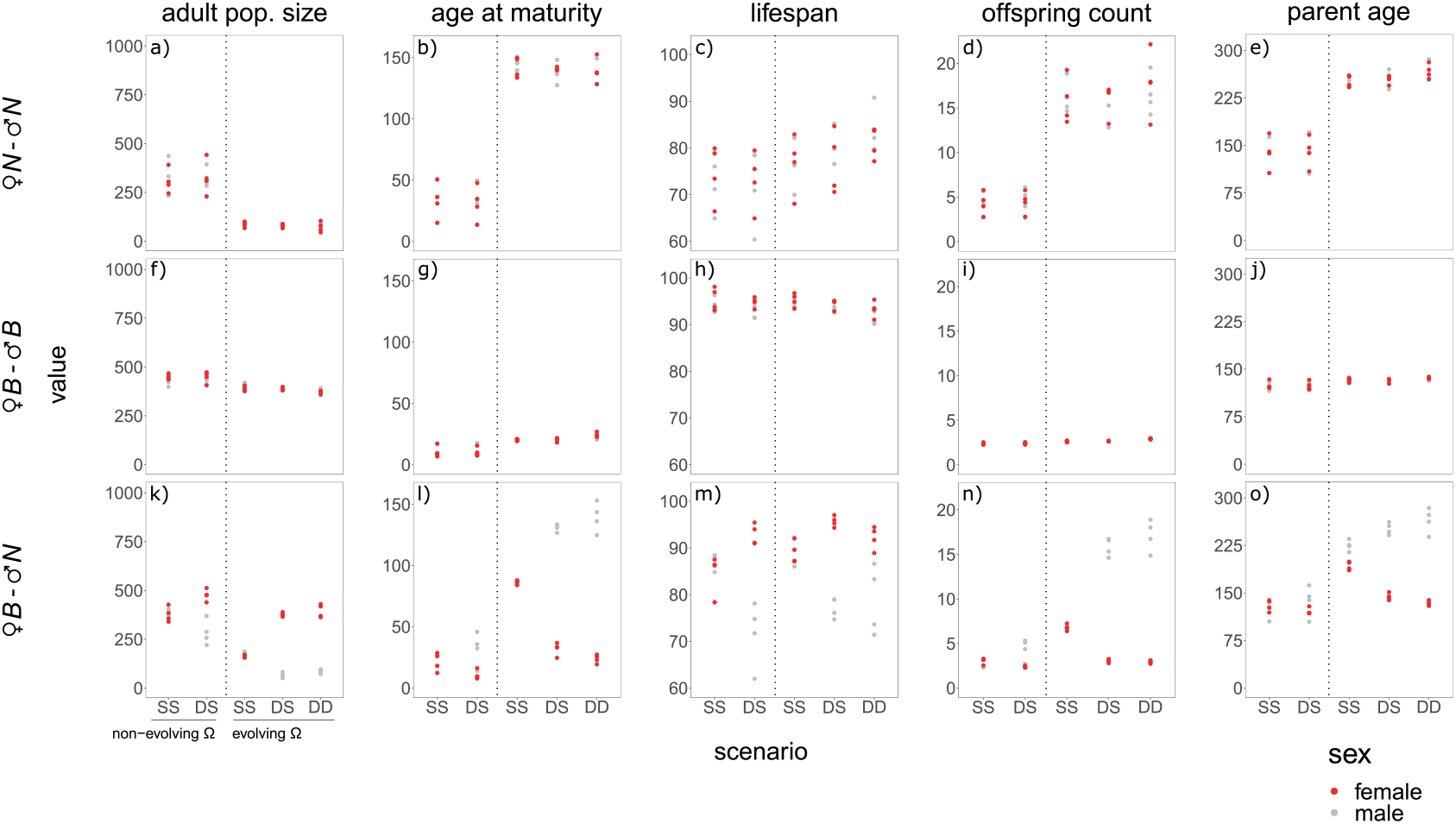
Life history trait evolution. Rows depict different reproductive competition scenarios as described in Fig.2. Columns show adult population size (number of mature individuals), mean age at maturity, average lifespan, offspring count (mean number of offspring of all the mature individuals), and parental age, with values in the y-axis of each panel. Adult population size is measured in the final time step and all the other traits are averaged across the population (mature individuals only, except for the lifespan) for the last 1000 time steps. Red gives the mean for females, grey for males. X-axis refer to different assumptions of sex-specific expression and ontogeny management evolution: **SS** when size and ontogeny management Ω traits are **S** hared between males and females, **DS** when size is **D** imorphic and Ω is **S**hared, **DD** when both are **D**imorphic; non-evolving Ω on the left of the dashed lines and evolving on the right, as indicated in panel k. Life histories reflected the differences in evolved target mature size (Fig.2). All parameters as in Fig.2.

#### Sex differences in reproductive scenario imply sexual conflict and can lead to sexual size dimorphism

The above results show that body sizes evolve very differently depending on whether the organism is selected to optimize its budget or whether selection instead simply favours the largest individual. This suggests potential conflict if selection is budget-based in one sex and size-based in the other, yet both have to express the same growth pattern and maturity size (a scenario with unresolved sexual conflict). In these situations we found size selection to override budget selection, visible in that *w_M_* = 1 and differing selection (Fig.2k, m, p, and r) yields evolutionary trajectories that are much more similar to the top row in Fig.2 than to the budget-based second row. In other words, both sexes increased their body size even if budget considerations made this suboptimal for one of the sexes.

When constraints on evolving sexual size dimorphism were lifted (*w_M_* = 0; Fig.2l, n, o, q, s, and t), sexual size dimorphism evolved in the expected direction: the sex that was selected for size evolved to be the larger sex. Sex differences in size were more pronounced when Ω was evolvable, with sexes differing nearly two orders of magnitude in size. Sex-specificity of ontogeny management made the sexes evolve towards different optima for some of the trait values (Supplementary Information, Fig.S2), but this had little effect on overall body sizes reached (Fig.2n and s compared to Fig.2o and t, respectively). The emerging sex differences in ontogenetic management Ω, when dimorphism in its composite traits was permitted to evolve, suggests the potential for sexual conflict can extend to quite fundamental ontogenetic traits such as thresholds of cell-level damage that trigger apoptosis (Supplementary Information, Fig.S2).

#### The initial strategies differ in their response to selection if not evolvable, but not in their evolvability

We used four lineages, all initially optimized for their ancestral size but differing in the combinations of traits that comprise their ontogenetic management strategy Ω. When all lineages were constrained to use their initial ontogenetic management strategy, some showed better preadaptations for size increases than others: Ω_4_ (purple diamond in Fig.2) in particular only permitted a very small size increase. Ω_2_ and Ω_3_ (black circle and blue triangle, Fig.2, respectively) permitted larger size increases than the other strategies, with Ω_3_ (red square) showing situation-specific constraints: if large size was selected for in both sexes or when sexual size dimorphism was permitted, it reached sizes close to Ω_2_. As a whole, Ω_2_ was the most prone to yield large bodies over evolutionary time, even in situations where all other lineages shrunk in size (Fig.2f-g, as well as Fig.2l and q for the budget-selected sex).

Making Ω evolvable (Fig.2c-e, h-j, m-o, and r-t) yielded a much more consistent body size response between the four independent lineages. Therefore, varying the potential rate of evolutionary change (for which we present two extreme options: absent and fast) in ontogenetic traits can shift body size responses towards higher repeatability and predictability, as a result of lifting the initially present constraints where ontogenetic management of small bodies is not suited to be scaled up to the development of larger bodies.

#### Life history traits scale with body size

The evolutionary trajectories of body sizes, described above, were accompanied by variation in life histories. Ontogenetic management evolution helped lineages to become large-bodied, and this reduced adult population sizes (Fig.3a). Large body sizes also led to an increased age at maturity despite the shortened expected lifespan, with mean lifespan falling below the age at maturity in some cases (Fig.3b-c), indicative of large juvenile mortality which helps to explain small adult populations in these cases. When the relatively few mature individuals of large-bodied species were responsible for all offspring produced, the offspring count (per mature individual; Fig.3d) and mean parental age (a measure of generation time; Fig.3e) increased. Sexual dimorphism in life histories reflected the evolution of sexual size dimorphism; when sexes evolved different mature sizes, they also varied in life history traits (Fig.3k-o).

#### Death causes depend on body size and can differ between the sexes

Across all scenarios and for both sexes, extrinsic mortality was the most common cause of death (Fig.4). Lineages suffered an increased risk of developmental failures (higher percent of deaths caused by stem cell depletion before maturation) if they evolved large bodies without coevolving adaptations in ontogeny management (Fig.4a, b, k, and l). Coevolution of ontogenetic strategies alleviated the problem of mortality caused by poor stem cell management and permitted attaining large body sizes, but this came with higher cancer mortality (Fig.4c-e, m-o). It is notable that the former leads to death before maturity and thus zero fitness, while a cancer-prone organism can reproduce (potentially multiple times) before succumbing to cancer; our result that evolving ontogenies ‘favour’ cancer over stem cell mismanagement can be understood in this light.

**Figure 4:**
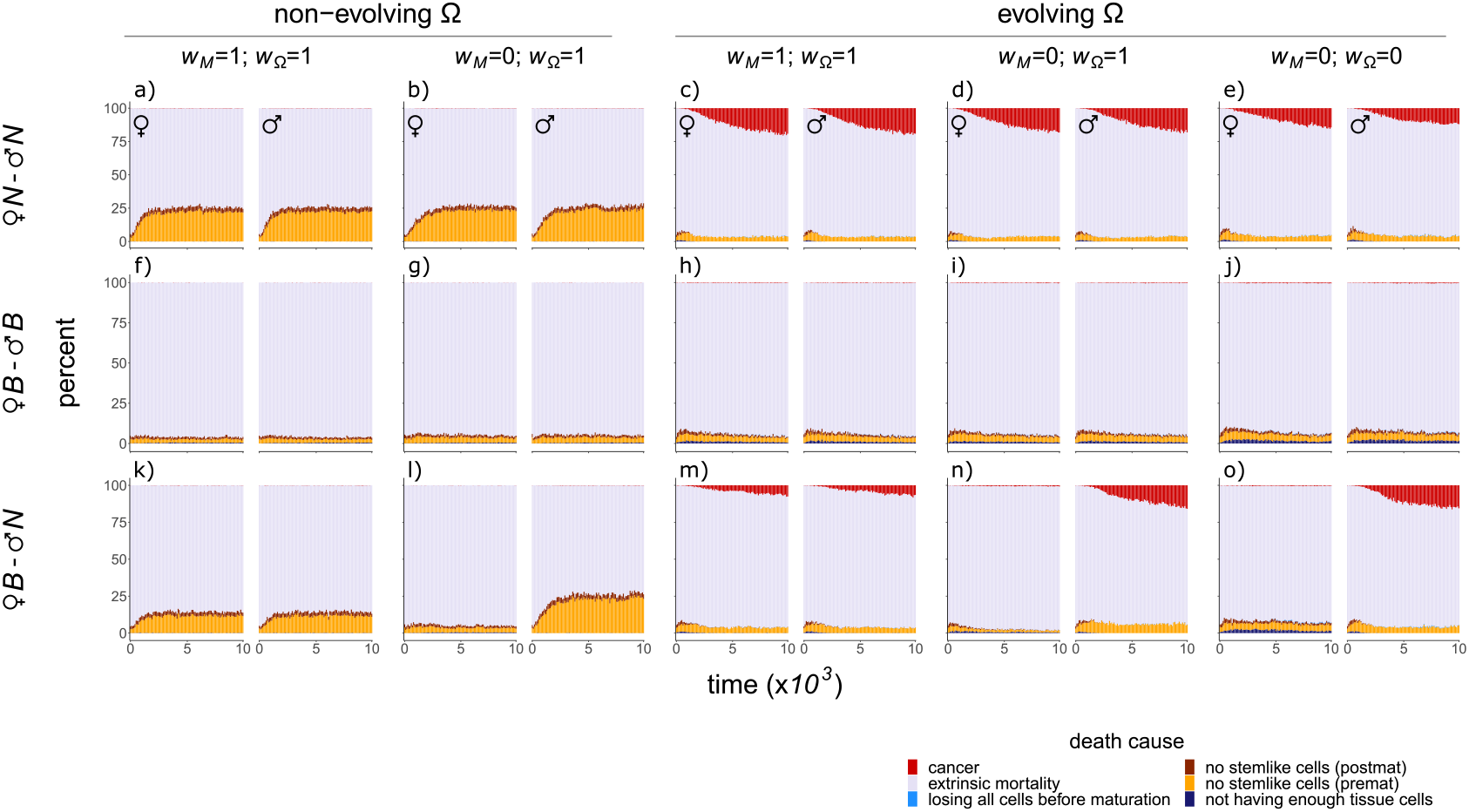
Death causes. Time is binned into intervals of 100 time steps (x-axis) and y-axis shows the percent of different death causes (distinguished with colours as in the figure legend) in each time bin, for all four initial ontogenetic strategies pooled together. Females on the left and males on the right in each panel, as indicated. Different reproductive scenarios, evolvability of ontogeny management Ω, and sex-specificity of body size (*w_M_*) and ontogenetic strategy (*w*_Ω_) depicted as described in Fig.2. Selection for an increased size and resulting body size evolution led to increased developmental failures (a, b, k, and j) or an increased incidence of cancer (c-e and m-o), depending on the scenario. Sexual size dimorphism resulted in differences between the sexes in terms of death causes (l, n, and o). All parameters as in Fig.2.

Differences in reproductive competition with sexually dimorphic body sizes (*w_M_* = 0, Fig.2) led the two sexes to differ in causes of mortality Fig.4l, n, and o). The larger sex suffered more from developmental issues (Fig.4l) if ontogenetic management did not coevolve with body sizes, and had a higher proportion of lives ending through cancer (Fig.4n-o) if ontogenetic management was permitted to coevolve with body size. Interestingly, the higher cancer risk of the larger sex (males in Fig.4) emerged whether or not the ontogenetic management strategy itself was permitted to evolve to be sex-specific (Fig.4n-o).

Expressing the same ontogenetic management strategy with the larger sex seemed to protect the smaller sex from failures in development to some extent (Fig.4n), compared to the other scenarios. When two sexes diverged in size but were constrained to the same genetic architecture for ontogeny management, Ω seemed to adapt to the needs of the bigger sex. This suggests that larger organisms require a more ‘specialized’ ontogeny management than the smaller ones. As a ‘side effect’, using a large-adapted ontogenetic strategy can protect smaller organisms from some mortality hazards during development (discussed in detail in Supplementary Information).

#### Cancer risk constrains body size evolution

Longer simulations did not make body size evolution reach a clear upper limit when ontogenetic management strategy was allowed to coevolve with size (the potentially open-ended scenario for size evolution, as shown above). However, these cases show a slowing down of the rate of increase in target mature size (Supplementary Information, Fig.S5). The fact that extrinsic mortality was the most prevalent death cause (Fig.4) and that average lifespan fell below the age at maturity in the largest lineages (Fig.3b-c) suggests a prominent role for extrinsic mortality selecting against ever longer maturation times, that even larger bodies than the observed ones would associate with. Simultaneously, cancer emerged as an important mortality risk in the largest lineages (Fig.4c-e, m-o).

To be able to estimate the role of cancer in hindering size increases, we ran additional simulations that completely eliminated cancer risk by not permitting any oncogenic mutations that lead to cell damage. These no-cancer simulations were conducted for two scenarios: (1) reproductive success is size-dependent in both sexes, (2) it is size-dependent in males while follows the budget selection model in females. Eliminating cancer risk allowed most lineages to evolve an increased target mature size (Fig.5), and in scenarios where body sizes evolved to be large under cancer risk, the further increase permitted by the no-cancer assumption was remarkable: in Fig.5, the points near log_10_ *M* =3 associate with log_10_ *M*_cancer−free_ ≈ 3.5, a more than threefold difference (0.5 difference on a log_10_ scale is 10^0.5^ = 3.16). The largest difference we found was between *M* =924 and *M*_cancer−free_=3603 (mean size of females with Ω_4_, when both sexes are size-selected, Ω is evolvable, and both size and Ω are sex-specific, averaged for the last 1000 time steps of the simulations). Overall, cancer plays a more important role in limiting size increases in large than in small organisms. Our simulations are in line with our analytical results where, all else being equal, selection against an equivalent proportional increase became stronger with increasing body size (Fig.1).

**Figure 5:**
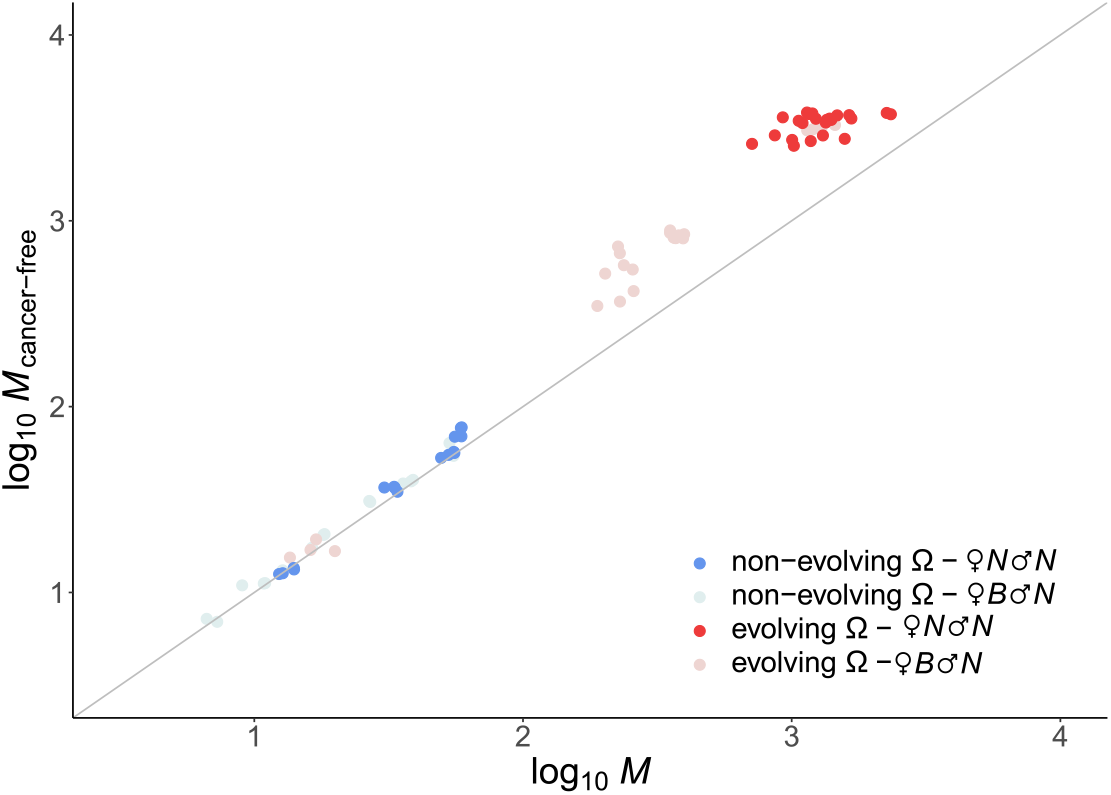
The effect of cancer risk on body size evolution. The evolved body size without cancer risk *M*_cancer*−*free_ (y-axis), plotted against the corresponding evolved size at the simulations with cancer risk (log_10_ *M* ; x-axis), with the diagonal line indicating *M*_cancer−free_ = *M*. Body sizes are averaged across the final 1000 time steps of the simulations. Colours depict evolvability of ontogenetic strategy and different reproductive scenarios, as indicated. Each point corresponds to a different combination of *w_M_*, *w*_Ω_ (as in Fig.2a-e and k-o) and sex. Overall, lineages tended to evolve larger body sizes when cancer-free than under cancer risk. Size increases were greater for the scenarios that resulted in large-sized lineages under cancer risk compared to those ended with small sizes. All parameters except for oncogenic mutation rate (which is *c*=0 for *M*_cancer−free_) as in Fig.2.

## Discussion

Our model builds on the previous work on body size evolution under cancer risk (Kokko and Hochberg, 2015) and the trade-off between cancer risk and reproductive success (Boddy et al., 2015), expanding them in two directions: by providing a firmer analytical footing of the basic expectation that the dangers of cancer increase with body size, and by adding differences between sexes and multiple ways in which a developing body may fail early or late in life.

Our two complementary models show that cancer risk can act as a constraint to body size evolution. The first model derives an analytical expression that indicates stronger selection against a proportional size increase in large-sized organisms, if cancer risk is the sole selective force. This model, by virtue of ignoring all reasons to be large, can be used to quantify the cost of further increases in size; should the benefits be constant, the increasing costs will at some point halt further increases. Our second model, using simulations, relaxes various restrictive assumptions of the analytical model. It includes explicit, and potentially sex-specific, ways how size can aid in reproductive competition. The various contrasts of the simulation shows that, (i) differences in life history (e.g. adult mortality or the probability of reaching the age at maturity) and population size can have their roots in ontogenetic management, forming a sufficient reason behind sex-specific life history traits acting alongside ecology, (ii) sexual conflict can play a role in cancer risk, creating sex-biased cancer incidence and potentially limiting the degree of sexual dimorphism, (iii) evolutionary increases in body sizes are limited, unless the lineage can lift this constraint by concurrent adaptation of its ontogeny management to its new body size, and (iv) if ontogenetic adaptation is slow or impossible, lineages can differ idiosyncratically in how pre-adapted their existing developmental strategies are for size increases (should there be selection for large size). Below, we discuss each of these results in detail.

First, in our model, evolving variation in body sizes resulted in differences in not only cancer incidence, but also impacted the population sizes of mature individuals and their life histories. Large organisms had small adult population sizes, reflecting a pervasive pattern in nature (Damuth, 1981; Juanes, 1986). Intriguingly, we did not explicitly model underlying ecological and energetic factors that might lead to small population densities in large organisms. Instead, this pattern emerged solely from the challenges of trying to reach a large adult size under constant size-independent extrinsic mortality. In reality, extrinsic mortality tends to decrease with increasing body size (McCarthy et al., 2008) and larger animals typically have a longer average lifespan (Blueweiss et al., 1978; Healy et al., 2014), allowing their life histories to explore options where maturation is delayed relative to small organisms. On the other hand, larger bodies require more resources to grow and maintain than smaller ones (Savage et al., 2004), which is typically argued to keep the population sizes of large organisms low (Peters, 1986). In turn, small population sizes can increase extinction risk both stochastically (Flather et al., 2011) and by slowing the rate of evolution and decreasing the adaptive potential in changing environments (Willi et al., 2006; Lanfear et al., 2014), potentially contributing to the apparent rarity of gigantism in extant taxa (Clauset and Erwin, 2008). Our result does not negate the importance of any of these ecological or macroevolutionary reasons behind population size, but it adds a reminder that time required to grow to mature size represents a greater challenge to large than to small organisms.

Our first result also relates to sexual dimorphism, as life history differences arose not only between different lineages, but also between the sexes if reproductive competition was settled differently in males and females. This led to biased adult sex ratios when size was sexually dimorphic, as well as shorter lifespan in the larger sex, in line with the literature (e.g. Ancona et al., 2020). Again, it is notable that these effects arose in our model without explicit assumption about underlying ecological and energetic factors that might lead to sexually dimorphic mortality: we did not assume that the large sex requires more resources to survive (Kokko and Brooks, 2003) or adopts more risk-prone behaviours (Kraus et al., 2008) or invests less in lifespan-prolonging traits such as immunocompetence (Foo et al., 2017; Kelly et al., 2018). Much of the effect in our model arose by growth taking time, and if growth cannot be sped up, a longer time to maturity is needed, giving extrinsic mortality time to act.

Second, the larger organisms (and the larger sex) also had their life more often cut short by cancer. Within a species, cancer incidence seems to increase with size (Albanes et al., 1988; Green et al., 2011; Fleming et al., 2011; Nunney, 2013, 2018). With the exception of humans and domestic dogs, relatively little is known about sexual dimorphism in cancer (outside obvious cases such as cancers in reproductive tissues that can only occur in one sex). In humans and domestic dogs, tissues that occur in both sexes appear to typically have higher cancer incidence in the larger sex (males; Ashley, 1969; Frank, 2007; Radkiewicz et al., 2017; Nunney, 2018), with size dimorphism at least partially explaining the observed differences (Walter et al., 2013; Nunney, 2018). Recent studies also point to potential sex differences in genetic and molecular trajectories of cancer (Li et al., 2020), in addition to the role of sex hormones in both promoting or preventing cancer (Kim et al., 2018). Our results, together with the available evidence, suggest that there is potential sexual conflict over ontogenetic management when cancer defences are ideally expressed to a different degree depending on sex-specific body size and life history requirements. Sexual conflict can also limit dimorphism itself via a similar route: in our model, when sexes differed in the type of reproductive competition, the large sex did not reach as large body size as when both sexes experienced selection to be large.

Third, size-selection in both sexes allowed lineages to increase in size without the constraints of sexual conflict. However, size evolution quickly reached an upper limit at relatively small sizes when ontogeny management was not evolvable. Ontogenetic management evolution permits coadaptation to novel body sizes, but even then, cancer risk remained a factor that keeps organisms smaller: our alternative simulations where cancer is impossible led to manyfold larger body sizes in an equivalent evolutionary time, with this difference being marked for the largest lineages considered. These results link to empirical evidence and theoretical studies highlighting the role of cancer-related and ontogenetic adaptations (Abegglen et al., 2015; Caulin et al., 2015; Kokko and Hochberg, 2015; Sulak et al., 2016; Tollis et al., 2019; Martinez et al., 2020; Vazquez and Lynch, 2020; Erten and Kokko, 2020; Nunney, 2020). In our model, we either permitted or did not permit ontogenetic adaptation to occur throughout the simulation; in reality, genetic changes for higher cancer suppression may in some cases have preceded size increases (Vazquez et al., 2018). This links to our finding within scenarios that lack continual adaptation: they vary in the range of sizes that a lineage can adaptively explore, based on idiosyncratic features of past adaptation to their current body size. Ontogenetic innovations might have been essential for permitting lineages to explore larger size ranges (e.g. increasing the number of oncogenic steps, Nunney, 2020), whereas adaptations in gene regulatory networks might have played a crucial role in enabling increased organismal complexity under cancer risk (Domazet-Lošo and Tautz, 2010; Trigos et al., 2019).

With our fourth result, we show that the ontogenetic management of larger bodies is challenging: the ‘rules’ that are sufficient to build a small body can fail if simply scaled up to bodies that require more cell divisions. In a broader, macroevolutionary perspective, there is little consensus about the underlying mechanisms for Cope’s rule (Stanley, 1973; Kingsolver and Pfennig, 2004; Heim et al., 2015; Gotanda et al., 2015). Apparent stasis together with occasional bouts of size increases may be linked to ‘innovations’ that allow large sizes (Payne et al., 2009), though taxon-specific factors (e.g. conflicting selection between larval and adult fitness components, Waller and Svensson, 2017) may provide alternative reasons to stay at a constant size despite an expectation that selection should favour increasing sizes. Increases in size seem, overall, harder to achieve than decreases (Evans et al., 2012), a result that is also borne out by our earlier work on size-specific cancer risk (Erten and Kokko, 2020).

Our current model also reiterates the fact that lineages differ idiosyncratically in how much they could respond to selection, based on existing ontogenetic rules. Even the most responsive lineages remained small, when body size was the only evolving trait, compared with cases where ontogenetic strategies were allowed to coevolve with body size (both experiencing abundant mutations). We suspect that reality could be between these two extremes: all lineages probably are somewhat able to respond to the selective pressure to increase cancer defences and, generally, organize their development and cell population maintenance to the challenges that occur at large size, but to also potentially differ in how easily the relevant innovations or adjustments arise. Intriguing evidence for the latter effect exists in the form of pleiotropic effects with clear potential to constrain the accessible adaptive landscape of cancer defences.

Looking for interspecific trends in cancer incidence has been instrumental for discovering novel mechanisms such as high p53-mediated apoptosis response in African elephant cells (Abegglen et al., 2015) or early contact inhibition in naked mole rat fibroblasts (Seluanov et al., 2018). Our study points out additional uses for examining outliers in comparative cancer data (both in terms of higher than expected and lower than expected cancer rates across species): it could help us not only find repeatable (across taxa) or taxon-specific mechanisms that offer protection from cancer, but can also shed light into how cancer risk might constrain the permissible range of body sizes that lineages can explore over evolutionary time.

## Acknowledgements

This work was supported by the university research priority program (URPP) “Evolution in Action” of the University of Zurich, as well as NIH grant U54 CA217376.

## Code availability

MATLAB code for the simulations is publicly available in GitHub: https://github.com/yagmurerten/Canstraint.

## Data availability

Mathematica notebooks used in analyeses, data derived from raw simulation data, and scripts used for processing and visualization are deposited in figshare, available at https://doi.org/10.6084/m9.figshare.13537295.

## Author contributions

EYE and HK conceived the study and developed the model, EYE performed the simulations and visualizations, EYE and HK wrote the paper.

