## Supplementary Information for "Cancer risk and sexual conflict as constraints to body size evolution"

### Analytical model with other body size correlations

In deriving Fig.1 in the main text, we assumed that body size affects lifespan via cancer risk with no other implications on fitness. Real relationships between body size and fitness incorporate various life history traits (Blueweiss et al., 1978), and it is not a priori clear whether each of them would make the negative effects of body size on selection stronger or mitigate against them. For example, we assumed that the probability of reaching maturity, $p_{\mathrm{mat}}$, does not change with body size. In reality, the requirement to grow to a mature size presents, all else being equal, a bigger challenge in a larger organism (the larger sex suffering higher juvenile mortality, Ancona et al., 2020), which would suggest a decline of $p_{\mathrm{mat}}$ with body size, translating into even stronger selection against size increases in large organisms than depicted in Fig.1. In a lineage evolving towards larger size, however, large parents may also explore a larger range of offspring sizes, including creating larger eggs (or neonates) than what a small-bodied organism could even in principle achieve. Offspring and adult sizes indeed covary in nature (Falster et al., 2008), which means that $p_{\mathrm{mat}}$ could conceivably increase with body size (Ronget et al., 2018). Our model does not directly comment on the likelihood of each possible relationship between $p_{\mathrm{mat}}$ and $N$, as size adjustments of offspring produced come with a complex set of questions such as the size-number trade-off in brood size evolution (present at least since the Cambrian, Ou et al., 2020). Our analytical model, considering cancer risk in isolation from other aspects of size dependence, shows the decline of $s$ with $N$ to reach strong values, thus it would require considerable (though possible) increases in $p_{\mathrm{mat}}$ with $N$, or other aspects of life history, to overcome selection to be small.

Other factors that may intensify selection against further size increases for large organisms relate to the fact that attaining a large size either takes more time or has to result from a faster rate of cell divisions. In the absence of compensatory mechanisms, the former means a lower $p_{\mathrm{mat}}$, whereas the latter could translate into a higher $k$ if fast growth can only be achieved by means of reduced 'quality control' such as DNA repair (Mangel and Munch (2005); for discussion of evidence see Ricklefs (2006) for speed of embryonic development, Bonisoli-Alquati et al. (2018) on associations between development time and DNA damage, and Péron et al. (2010) and Cooper and Kruuk (2018) for effect of early developmental conditions on senescence; these analyses, however, are somewhat tangentially relevant to our central question as they test neither for body size effects nor cancer).

Nevertheless, there are also selective advantages of being large such as low extrinsic mortality (Savage et al., 2004; McCarthy et al., 2008), Below, we assume the extrinsic mortality rate scales with body size as $\mu = e^{-0.5}N^{-0.25}$ (following the relationship for mammals in McCarthy et al., 2008). This yields a case where selection on size switches from positive at small body sizes to negative at large ones, i.e. stabilizing selection for an intermediate size (Fig.S1). In other words, including allometric benefits of a large body size (in this case a reduction of extrinsic mortality with size) can change the results from a uniform decline in $s$ with $N$ to one where smallest organisms are selected to increase in size, but it does not destroy the argument that cancer forms a constraint that prevents unbounded size increases.

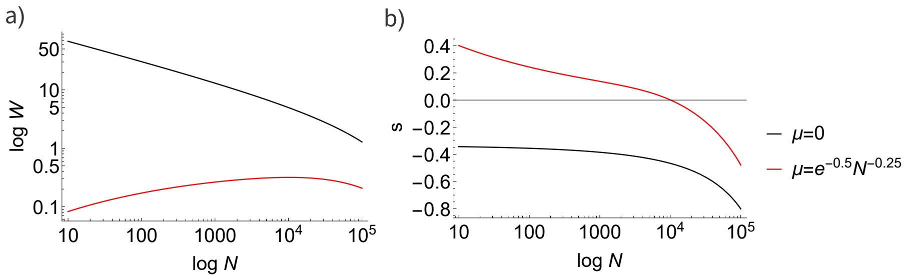


Figure S1: Fitness and selection when extrinsic mortality scales with body size. (a) Fitness ($W$) across body sizes ($N$), both in log-scale, following Eq.2. (b) Selection $s$, defined as elasticity (proportional change in fitness with a proportional change in body size) according to Eq.3. Both are plotted against $\log N$ and evaluated for $t_{\text{mat}}$=2, $p_{\text{mat}}$=1, $k$=0.005, and $n$=3; with $E(t) = 1-e^{\mu}$, where $\mu = 0$ corresponds to our results in the main text (black), and $\mu = e^{-0.5}N^{-0.25}$ is when we assume a scaling relationship between size and extrinsic mortality (red). With this scaling relationship, highest fitness is attained at intermediate body sizes and there is stabilizing selection for an intermediate size (b).

### Additional results of the sexual model

In this section, we present additional results of our sexual model: 1) the evolution of individual traits that make part of ontogeny management $\Omega$, when the latter is evolvable, and the trajectory of body size evolution 2) in the scenarios with partially sex-specific size and $\Omega$ expression, 3) without cancer risk, 4) when we ran our sexual model for more time steps than in the main text.

#### Evolution of ontogenetic management strategy

In the main text, we show that lineages can attain markedly larger sizes than their initial mature size if ontogeny management $\Omega$ is evolvable (under scenarios with selection for size), suggesting that large size requires specific $\Omega$ adaptations. Indeed, some of the individual traits that make part of $\Omega$ seem to have different optima in large-sized organisms (when both sexes are size-selected; Fig.S2a-g) compared to small-sized ones (when both sexes are budget-selected; Fig.S2h-n). For instance, compared to small-sized lineages, large-sized ones typically had a higher threshold for DNA damage response ($A$; Fig.S2d, k, for large- and small-sized respectively), more differentiation levels ($T$; Fig.S2f, m), which themselves had lower relative division propensity compared to the previous level ($X$; Fig.S2g, n). Overall, the highly repeatable values under budget selection, i.e. the selective regime that led to small sizes (Fig.S2h-n), were replaced with much more idiosyncratic ontogenetic management strategies when evolving to be large (under size selection, Fig.S2a-g), visible as more variable outcomes in the latter case.

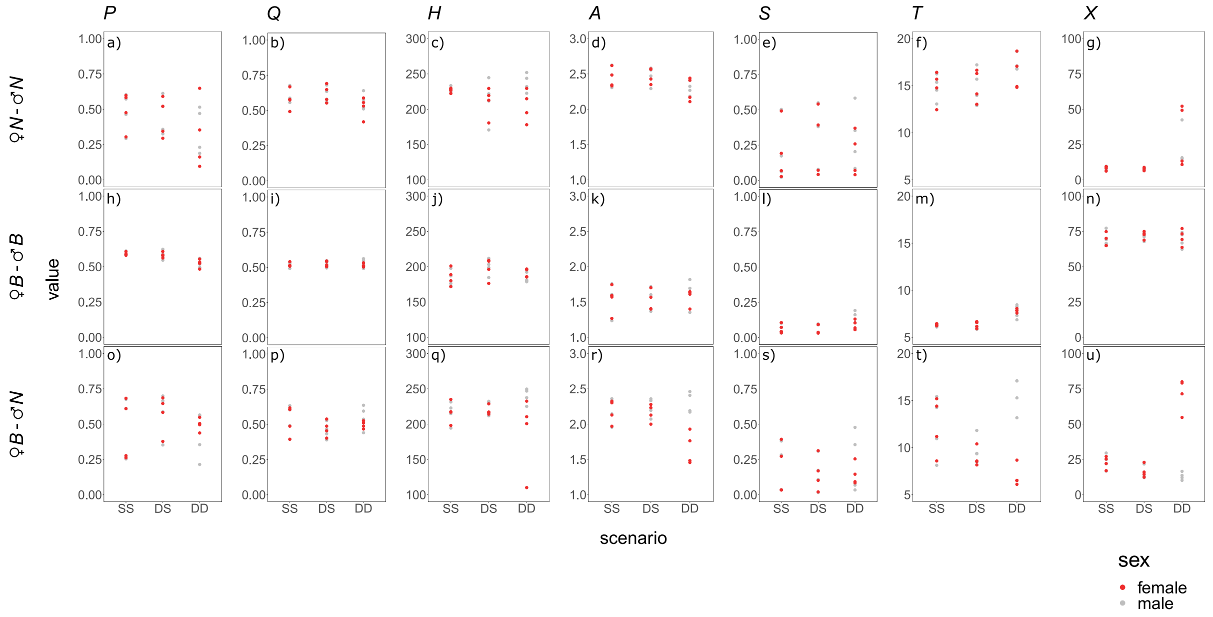


Figure S2: Final trait values comprising the evolving ontogenetic management strategy $\Omega$, averaged across the mature population for the last 1000 time steps, separately for females (red) and males (grey). The four replicates reflect evolution that started from four ancestral populations (see main text). Columns, from left to right: probability of asymmetric cell divisions, $P$; probability of differentiation in symmetric divisions, $Q$; maximum number of times a cell can divide, $H$; DNA damage response threshold, $A$; DNA damage response strength, $S$; number of differentiation levels from a stem cell to a terminally-differentiated cell, $T$; division propensity of a differentiation level compared to the previous one, $X$. Rows correspond to reproductive scenarios: size selection in both sexes ($N$-$N$), budget selection in both sexes ($B$-$B$), budget-selection in females and size-selection in males ($B$-$N$). The x-axis shows different sexual conflict scenarios: **SS** when size and ontogeny management $\Omega$ traits are **S**hared between males and females, **DS** when size is **D**imorphic and $\Omega$ is **S**hared, **DD** when both are **D**imorphic. Parameter values: extrinsic mortality $\mu$=0.01, cell turnover rate $\nu$=0.0001, oncogenic mutation rate $c$=0.01, number of oncogenic steps $n$=3. Some ontogenetic traits have size-specific optima ($A$, $T$, and $X$) and show sexual dimorphism when sexes differ in their reproductive scenario and can evolve different body sizes and ontogeny management (DD scenarios).

When the type of reproductive competition was sex-specific but sexes shared their ontogeny management and size expression, selection for size overrode the demands for budget optimization (Fig.2 in the main text). In these cases, $\Omega$ evolution resembled the scenarios in which both sexes were size-selected (e.g. $X$, SS scenario in Fig.S2g, n, u).

Allowing sex-specific expression of body size resulted in sexual size dimorphism, implying different optimal $\Omega$ trait combinations between the sexes, and therefore sexual conflict when $\Omega$ was not permitted to evolve sex specificity. Sexually antagonistic selection resulted in intermediate values for some traits (e.g. $T$, DS scenario in Fig.S2f, m, t) and the evolution towards the values optimal for the larger sex in others (e.g. $X$, DS scenario in Fig.S2g, n, u). These traits relate to differentiation strategies in terms of tissue structure; high $T$ translates into (1) more divisions until terminal differentiation, and, combined with a high threshold for damage response $A$, but otherwise similar traits ($P$, $Q$, $H$ and $S$, trait definitions given in Fig.S2), (2) more transient stem cells between the least differentiated level and the somatic tissue.

Using a tissue structure (i.e. $T$ and $X$) adapted to a large-sized body can explain the lower percent of stemlike cell related deaths in the small sex when $\Omega$ was shared (Fig.4n in the main text, females compared to all the other panels). Additionally, requiring more cell divisions until terminal differentiation can underlie a slightly higher age at maturity of the small sex when $\Omega$ was shared compared to when sexes can optimize their $\Omega$ separately (Fig.3l in the main text, DS compared to DD). These differences were also accompanied by a slightly longer lifespan (Fig.3m in the main text, DS compared to DD). However, a suboptimal strategy can incur fitness costs despite an increased lifespan (Maklakov and Lummaa, 2013), which is also visible in our model. Maintaining a higher number of stemlike cells (due to (2) above) lowers the rate of budget accumulation in our model ($b(t)$) and can translate into lower resources available for reproduction. Since competition was settled based on relative budgets in our model, the mean offspring count of mature individuals of the smaller sex was similar between shared and sex-specific $\Omega$ scenarios (Fig.3n in the main text). In a context where the absolute amount of resources affects reproduction (e.g. Naulleau and Bonnet, 1996), the small sex can suffer from a low reproductive success using a large-adapted $\Omega$, in line with our earlier results (Erten and Kokko, 2020).

#### Partially sex-biased expression

In the main text, we considered two extremes for sex-biased gene expression: target mature size $M$ and ontogeny management $\Omega$ could be either fully shared between the two sexes or completely sex-specific. Here, we explored the intermediate cases where we allow gene expression to be partially sex-specific. When reproductive scenarios differed between the sexes, a partially sex-specific $M$ expression resulted in a short period of transition until the body size became fully dimorphic in some lineages (as can be seen in e.g. Fig S3m and o). Otherwise, the results exhibited no qualitative difference compared to the corresponding scenarios (i.e. non-evolvable or evolvable $\Omega$) where we allowed dimorphism in the main text.

#### Body size evolution without cancer risk

In the main text, we reported the evolved target mature size without cancer risk in relation to the corresponding body size that evolved under cancer risk. To provide a more detailed view, here we show the trajectory of body size evolution over time without cancer risk (Fig.S4).

#### Simulations ran for longer time than in the main text

Due to computational limits, we ran our simulations for 10000 time steps in the main text. Here we show the body size evolution for a longer duration of 20000 time steps

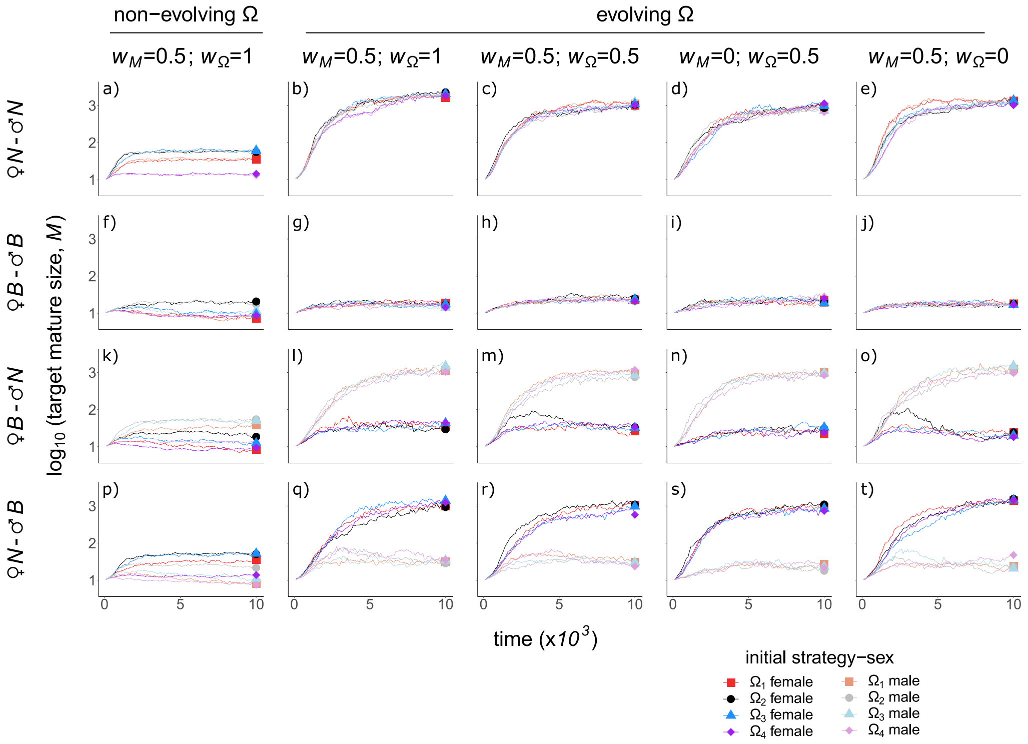


Figure S3: Body size evolution under partially sex specific trait expression. Columns indicate evolvability of the ontogenetic management as well as the sex-specificity of body size ($w_M$) and ontogeny management ($w_\Omega$); values of 1 indicate that sexes express shared alleles, 0 completely sex-specific expression, and 0.5 that shared and sex-specific alleles weigh equally in phenotype determination. Rows: reproductive scenarios as in Fig.S2, with an additional row showing size-selection in females and budget-selection in males ($N$-$B$). Other parameters as in Fig.S2. In each panel, x-axis: time steps in the simulation; y-axis: mean target tissue size in logarithmic scale ($\log_{10} M$), plotted for mature individuals in the population for every 100 time steps. Colours and dots show the initial strategy and sex as indicated.

(Fig.S5). Our results show that coevolving ontogenetic management with body size can sustain increased sizes, though with a tendency to slow down over time.

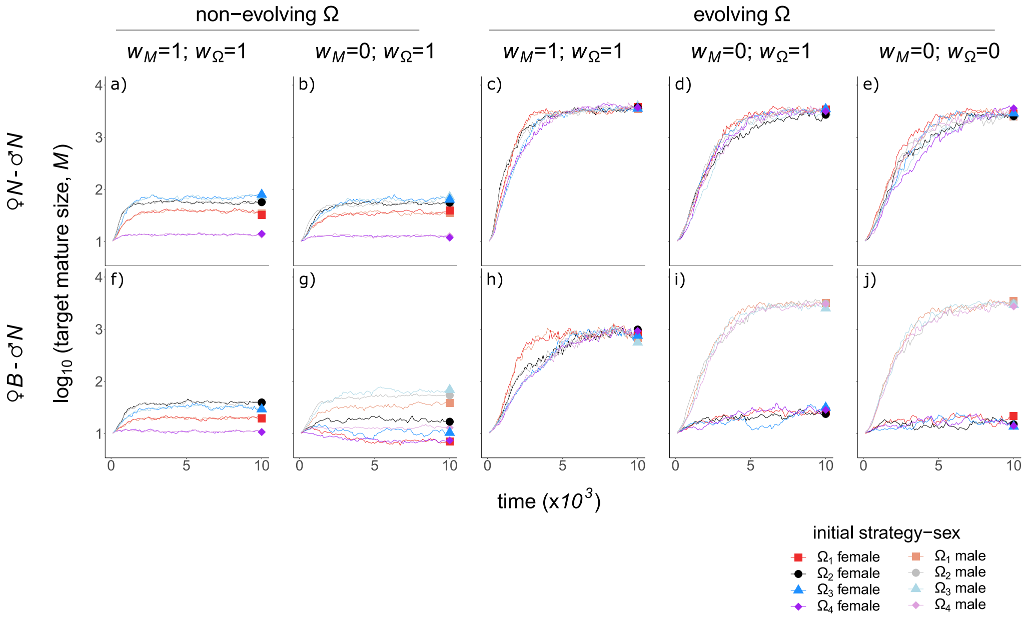


Figure S4: Body size evolution with no cancer risk. Cancer risk is set to zero ($c$=0), other figure details and parameters as described in Fig.S3. With no oncogenic mutations and an evolvable Ω, body size evolved to be larger than in the scenarios with cancer risk (Fig.2 in the main text).

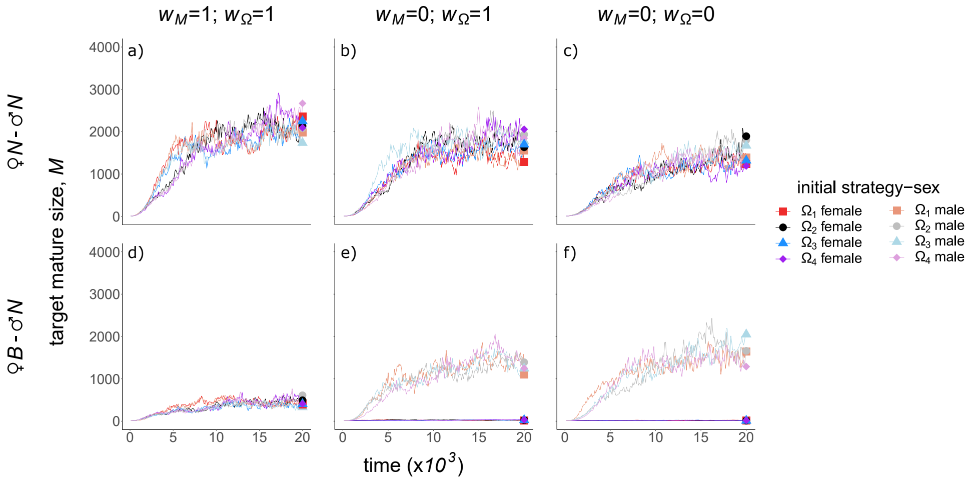


Figure S5: Body size evolution. The evolutionary trajectories of the body size change when simulations were ran for 20000 time steps, shown only for the scenarios with evolvable Ω. To make ongoing changes at large sizes clearer, we replace the logarithmic scale used elsewhere with a linear scale on the y axis. All other details are as described in Fig.S3. Body size increases continued over time, but became slower after the lineages evolved to a certain target mature size, depending on the scenario.
